# Benchmark of Differential Gene Expression Analysis Methods for Inter-species RNA-Seq Data using a Phylogenetic Simulation Framework

**DOI:** 10.1101/2022.01.21.476612

**Authors:** Paul Bastide, Charlotte Soneson, Olivier Lespinet, Mélina Gallopin

## Abstract

Inter-species RNA-Seq datasets are increasingly common, and have the potential to answer new questions on gene expression patterns across the evolution. Single species differential expression analysis is a now well studied problem, that benefits from sound statistical methods. Extensive reviews on biological or synthetic datasets have provided the community with a clear picture on the relative performances of the available tools in various settings. Such benchmarks are still missing in the inter-species gene expression context. In this work, we take a first step in this direction by developing and implementing a new simulation framework. This tool builds on both the RNA-Seq and the Phylogenetic Comparative Methods literatures to generate realistic count datasets, while taking into account the phylogenetic relationships between the samples. We illustrate the features of this new framework through a targeted simulation study, that reveals some of the strengths and weaknesses of both the classical and phylogenetic approaches for inter-species differential expression analysis. The tool has been integrated in the `R` package `compcodeR` freely available on `Bioconductor`.

## 1 Introduction

The study and analysis of gene expression differences across species is a long standing problem (King and Wilson, 1975). The development of microarray technologies led to the gathering of the first large scale and across species gene expression datasets, that allowed for the formulation and study of various hypotheses regarding the link between gene expression and evolution (Enard, 2002; Gilad et al., 2006; Khaitovich et al., 2004; Whitehead and Crawford, 2006). RNA-Sequencing technologies have changed the way to measure gene expression (Wang et al., 2009), making comparisons across several species easier, even for species with no reference genome available (Perry et al., 2012; Romero et al., 2012). As a compilation of a large number of datasets and atlases for gene expression of healthy wild-type individuals, the well maintained Bgee database (Bastian et al., 2021) is an important resource to ease the comparison of expression patterns across animal species.

Since changes in expression may underlie complex phenotypes, across species gene expression datasets can be used to test a wide range of evolutionary scenarios (Dunn et al., 2013; Romero et al., 2012). Tested hypotheses include for instance expression divergence (Gu, 2004); strength of expression conservation (Gu et al., 2019); coevolution of gene expression (Cope et al., 2020); test of the orthologous conjecture (Rogozin et al., 2014; Dunn et al., 2018); detection of the “phylogenetic signal” (Musser and Wagner, 2015); equality of within-species variance (Catalán et al., 2019); constant stabilizing selection, loss through drift, parallel or divergent selection (Stern and Crandall, 2018a,b); or detection of duplication-specific effects in expression evolution (Fukushima and Pollock, 2020).

In this review, we focus on the detection of change in gene expression levels across species, in a specific lineage or between different groups of species. This problem can be formalized as an inter-species differential expression analysis, and has been studied in various groups of organisms (Stern and Crandall, 2018b; Cáceres et al., 2003; Zheng-Bradley et al., 2010; Blake et al., 2018; Chen et al., 2019; Blake et al., 2020; Alam et al., 2020). For instance, difference in gene expression levels was found between mammalian lineages and birds (Brawand et al., 2011), across non-model primates species (Perry et al., 2012), between *Drosophila* species (Torres-Oliva et al., 2016) or *Heliconius* butterflies (Catalán et al., 2019). Note that the biological interpretation of changes in the level of expression of a gene across species is not easy (Romero et al., 2012). Shifts in gene expression across species could be molecular signatures of ecological adaptation, or associated with a directional selection scenario, or a relaxation of evolutionary constraints.

From a bioinformatic point of view, the comparison of RNA-Seq samples between multiple species requires, first, the detection of orthologous relationships between genes (Tatusov, 1997; Tekaia, 2016), second, the consideration of differences in genome mappability (Zhu et al., 2014) and, third, the adaptation of alignment and quantification pipelines (LoVerso and Cui, 2015; Chung et al., 2021). Multi-species alignments techniques have also been developed (Bradley et al., 2009; Brawand et al., 2011). In this review, we name orthologous genes (OG), or simply genes, the set of genes having orthologous relationship across species. Once the orthologous gene expression matrix has been created, the level of expression can be transformed into a discrete variable to detect presence versus absence of gene expression (Bastian et al., 2021). Other approaches perform separate differential expression for each species (Dunn et al., 2013; Kristiansson et al., 2013) or focus on pairwise comparisons only (Zhou et al., 2019; Chung et al., 2021). Direct comparisons of expression between species can be complicated by batch effects (Gilad and Mizrahi-Man, 2015), or potential confounding factors (Roux et al., 2015; Cope et al., 2020). Comparative gene expression studies should be carefully designed (Dunn et al., 2013; Romero et al., 2012; Chung et al., 2021). In this work, we focus our attention on genes having a one-to-one relationship across several species (more than two species). We consider the level of expression of genes as a quantitative trait evolving across several species, and we detect genes with a shift in the level of expression across species as performed in e.g. Brawand et al. (2011); Perry et al. (2012); Torres-Oliva et al. (2016); Stern and Crandall (2018a). Since no statistical method is clearly established to perform this detection across multiple species, we present in the next section a review of all strategies used in practice.

There are several well-established tools to simulate RNA-Seq count data in the classical, intra-species case (Dillies et al., 2013; Soneson and Delorenzi, 2013; Soneson, 2014), which allowed for the benchmark of many differential expression analysis models (Anders and Huber, 2010; Robinson and Oshlack, 2010; Law et al., 2014). Although some methodological questions remain open (Van den Berge et al., 2019), these extensive simulation studies helped setting good practices in terms of model choice or normalization methods in various intraspecies RNA-Seq settings. To our knowledge, there exists no extension of these frameworks to the inter-species setting. Simulation of gene expression across species has been performed using linear models and Gaussian variables (Rohlfs et al., 2014; Rohlfs and Nielsen, 2015; Gu et al., 2019), but without taking into account the specificity of RNA-Seq count data and without focusing on the detection of shifts across species. In this review, we propose a framework to simulate RNA-Seq data across species. We use this framework to compare different strategies to detect genes with a expression level shift across multiple species, and draw recommendations for inter-species gene expression comparison.

The paper is structured as follows: first, we describe the main methods used to perform differential analysis or shift detection across multiple species. We then explain our simulation method to generate synthetic inter-species RNA-Seq data using a Poisson log-normal model. Our simulation tool is integrated in the Bioconductor package `compcodeR`. Finally, a targeted simulation study, that draws its parameters from a recent inter-species RNA-Seq study (Stern and Crandall, 2018a), allows us to compare the current statistical methods, and propose some recommendations.

## 2 Review of methods used to compare level of expression across species

### 2.1 Setting and Notation

For the remainder of this work, *y_gi_* denotes the measured level of expression for gene *g*, 1 ≤ *g* ≤ *p*, and sample *i*, 1 ≤ *i* ≤ *n*. We assume that the species are partitioned into two groups *S*_1_ and *S*_2_, that depends on the biological question at hand. Each sample is associated to a species, and each species belongs to one of the two groups of interest.

Our goal is to detect genes with a shift in expression level across groups. To perform this test, we need to properly model the level of expression of gene *g* in sample *i*, taking into account the specificities of inter-species RNA-Seq data, that are multifold. Indeed, RNA-Seq data are counts, usually measured on a low number of samples. In addition, several technical biases affect the measured level of expression *y_gi_*, either gene-specific (such as heterogeneity of gene length and GC content across genes and samples), or sample-specific (such as heterogeneity in library size across samples). Finally, since the level of expression of a gene *g* is measured across several species, the phylogenetic relationships between species induce some correlations in the data. While, ideally, all these specificities should be taken into account in the statistical analysis, to our knowledge there exist no model that includes all these constraints in its hypotheses. Below, we present an overview of the three main strategies adopted to model inter-species RNA-Seq data, that each make different simplifying assumptions.

We denote by *m_i_* the sample specific normalization factor for sample *i*. Several approaches exist to compute this factor (Dillies et al., 2013), such as the Relative Log Expression (RLE) (Anders and Huber, 2010) method or the Trimmed Mean of M-values (TMM) (Robinson and Oshlack, 2010) method. We further denote by *ℓ_gi_* the length of the gene *g* in sample *i*, which need to be taken into account as a gene and sample specific normalisation factor.

All the methods described below rely on a (generalized) linear model. The design (or model) matrix **X** of the experiment defines the form of this model. For differential analysis, it contains at least a grouping information, specifying which biological replicate belongs to *S*_1_ or *S*_2_. It can include some covariates that might influence the gene expression, such as information about environmental or experimental conditions. The matrix **X** has *n* rows, and as many columns as the number of coefficients in the model.

### 2.2 Strategy 1: Generalized Linear Model on Raw Count Data

The first option to perform differential expression analysis across species is to use a generalized linear model based on the negative binomial distribution (Anders and Huber, 2010; Robinson and Oshlack, 2010), implemented in several `R` packages such as `DESeq2` or `edgeR`. In `DESeq2` (Love et al., 2014), the random variable modeling the raw level of expression *Y_gi_* of gene *g* in sample *i* is a negative binomial with expectation *μ_gi_* = *c_gi_q_gi_* and dispersion *α_g_*: *Y_gi_* ~ *NB*(*μ_gi_*,*α_g_*). The coefficient *c_gi_* is a sample and gene specific normalization factor that depends on the sample specific normalization factor *m_i_* and on the gene length *ℓ_gi_*. The parameter *q_gi_* is linked to the true level of expression of sample *i*, and includes the model design through the relationship log_2_(*q_gi_*) = **X**_*i*_.***θ***_*g*_, where **X**_*i*_. denotes the *i^th^* line of the design matrix **X**, and the vector of coefficients ***θ***_*g*_ contains the information on the log_2_ fold changes between the two groups of species for gene *g*.

This method properly models counts and is appropriate to analyse data with low sample size thanks to dispersion shrinkage (Anders and Huber, 2010; Robinson and Oshlack, 2010). Sample specific and gene specific technical biases are taken into account directly into the parametrization of the model. Unfortunately, to our knowledge, this model is not flexible enough to account for the correlation induced by the phylogenetic tree. For this reason, this model is usually used to perform pairwise comparison between species (Torres-Oliva et al., 2016).

### 2.3 Normalization and Transformations

As we will see below, instead of using a generalized linear model on raw count data, it is possible to use a simple linear model on normalized data. The normalization step is essential to transform count measurements into continuous values, and to unlock the use of linear models. The normalization should be designed to temper the sample and gene specific technical biases, as well as to render the data homoscedastic (i.e. with homogeneous variance across samples).

Three main normalization scores are used in the literature. They all rely on the normalized library size *M_i_* for sample *i*, defined as: *M_i_* = ∑*_g_y_gi_m_i_*, with *m_i_* the scaling normalization factor described above. The Count Per Million (CPM) score incorporates sample-specific normalization only: 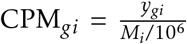. The Reads (or fragments) per kilobase per million mapped reads (RPKM) score incorporates an extra gene-specific normalization as follow: 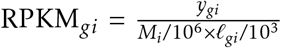 (Mortazavi et al., 2008). Another way to include the same gene-specific normalisation is to use the Transcripts per million (TPM) score: 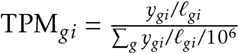 (Wagner et al., 2012). Compared to the RPKM, the TPM scores summed over all genes are equal to a constant (10^6^), which is a property that can be desirable in some settings (Musser and Wagner, 2015).

In addition to the normalization, an extra transformation is often needed to make the data behave closer to a homoscedastic Gaussian. Two transformations are widely used: the log_2_ transformation (Law et al., 2014) and the square root transformation (Musser and Wagner, 2015).

For inter-species differential expression analysis, the choice of the right normalization and transformation to perform is not clearly established. Some studies use the log_2_-transformed RPKM (Mortazavi et al., 2008; Brawand et al., 2011; Catalán et al., 2019) or CPM (Blake et al., 2018) scores. Other studies advocates for the use of the log_10_ (Chen et al., 2019) or square-root (Musser and Wagner, 2015; Stern and Crandall, 2018a) transformed TPM.

In the remainder of this work, *ỹ_gi_* denotes the normalized and transformed level of expression for gene *g* and sample *i*.

### 2.4 Strategy 2: Linear Model on Normalized Data

Assuming the data has been normalized and transformed properly, it can be modelled, for each gene *g*, using a simple linear regression:

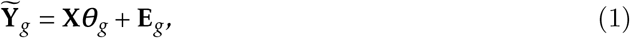

where 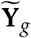 is the vector of the *n* normalized measurements for gene *g*, **E**_*g*_ is a vector of Gaussian independent and identically distributed residuals, and, as previously, **X** is the design matrix and ***θ***_*g*_ the associated vector of coefficients. This model is implemented in the popular `R` package `limma` (Smyth, 2004; Smyth et al., 2005), that uses an empirical Bayes moderated statistic to test whether the coefficient of ***θ***_*g*_ associated with the group segregation is significantly different from zero. This method is appropriate to analyze datasets with low sample size, but a large number of genes that are pooled in a hierarchical model to get a better estimation of the variance.

It can be applied directly to RNA-Seq data, normalized using the previous methods. If the data presents mean-variance trends, which is typically the case in classical intra-species RNA-Seq data due to the presence of a high number of highly variable small counts, this can be taken into account through a weighting method (`voom`), or through the direct inclusion of the trend in the hierarchical empirical Bayes model (the `trend` method) (Law et al., 2014).

This method does not take the phylogenetic correlations into account, and has been used to performed pairwise comparisons (Blake et al., 2018, 2020; Torres-Oliva et al., 2016). This model is flexible and can be extended to a linear mixed model that accounts for the correlation between replicates of the same species (Breschi et al., 2016), using the `duplicateCorrelation` function from `limma`. However, correlations between species, encoded by the phylogenetic tree, cannot be directly taken into account using this approach.

### 2.5 Phylogenetic Comparative Methods

The methods described above are tailored for RNA-Seq data, but they are not designed to deal with the correlations introduced in the measurements by the phylogenetic relationships between the samples in an inter-species analysis. In this section, we briefly introduce Phylogenetic Comparative Methods, that have precisely been developed to deal with these correlations, before demonstrating some of their uses in the RNA-Seq literature.

#### Phylogenetic Comparative Methods

Phylogenetic relationships are known to induce correlations between observed quantitative traits on several species (Felsenstein, 1985). The field of Phylogenetic Comparative Methods (PCMs) specializes in the comparative study of such phylogenetically related traits, and has been flowering over the last decades (see e.g. Harmon (2019) for a recent review). Conditionally on a phylogenetic tree that links a set of species, PCMs model the evolution of a quantitative trait as a stochastic process running along the branches of the tree (see Fig. 1). This generative model induces a multivariate Gaussian structure of the observed vector of traits across species, with a correlation structure that depends on the tree and on the chosen process. The values of the trait are only observed at the tips of the tree. The values at the root or at the internal nodes are unobserved and are modeled using latent variables.

**Figure 1:**
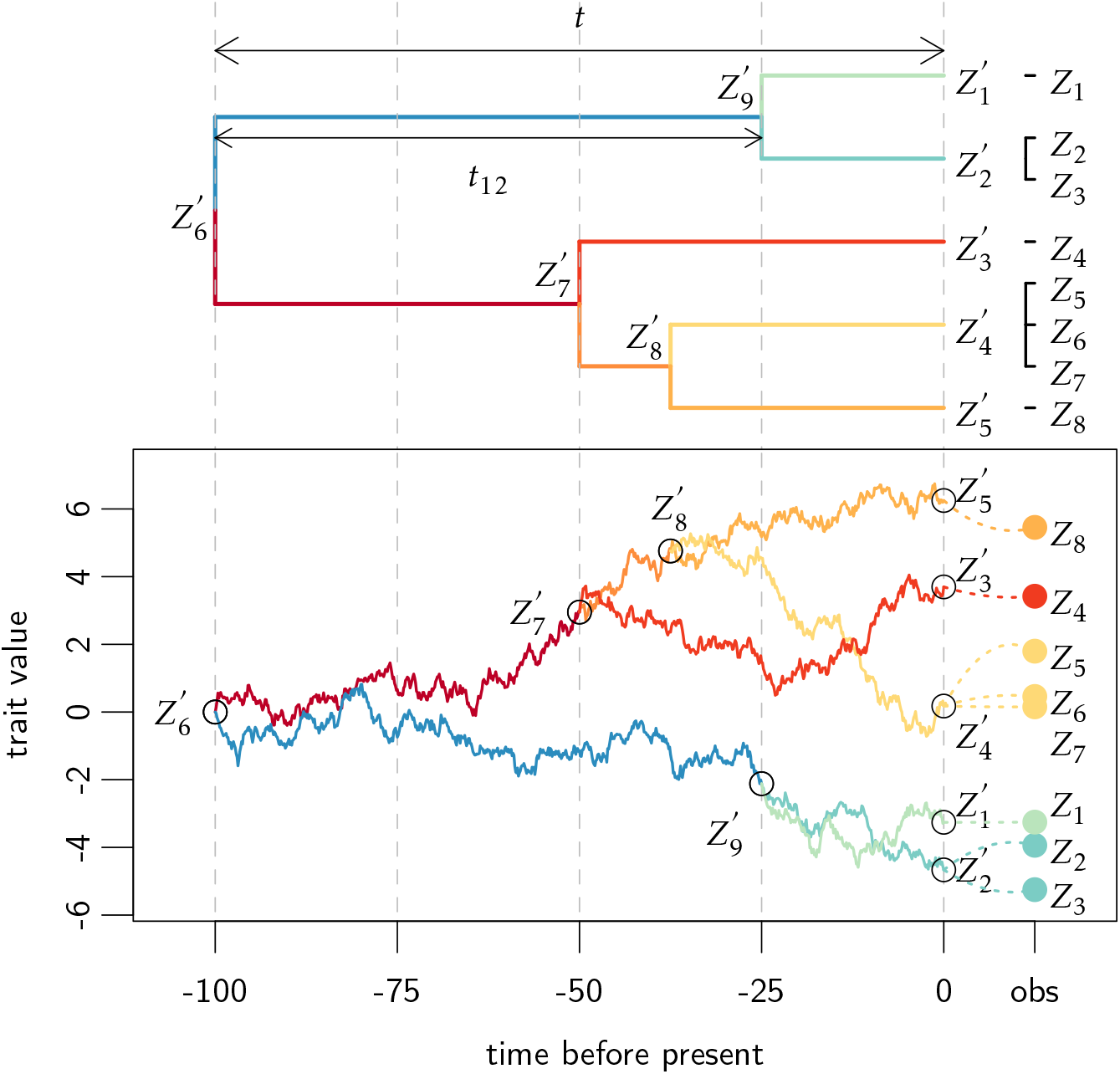
Realization of a Brownian Motion (BM) process (bottom), on a time calibrated ultrametric tree with total height *t* = 100 (top), with replicates and within-species variation. The BM process on the tree controls the distribution of the internal nodes, including ancestral nodes 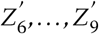, and latent tip traits 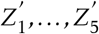. The ancestral root value of the BM is *μ* = 0, and its variance is 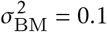, so that the latent (unobserved) tip trait variance is 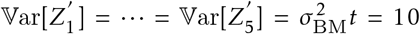. The covariance of the latent tips trait is proportional to their time of shared evolution, for instance 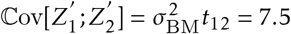. Replicated measurements are added on the tree as tips with zero branch lengths (top), with an extra variance of *s*^2^ = 0.5. For instance, *Z*_2_ and *Z*_3_ are replicates of the latent tip 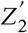, and their conditional distribution is Gaussian with expectation 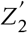 and variance *s*^2^. The total sample traits variance is hence given by 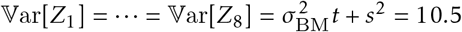, and the sample traits covariance is given by the tree structure, for instance 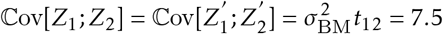, and 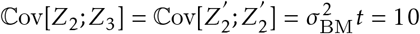. Note that on this figure, latent internal nodes (internal and external) are numbered from 1 to 9, and observations are numbered from 1 to 8, but these set of indices are distinct. For instance, *Z*_1_ is indeed an observation of 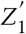, but *Z*_4_ is an observation of 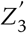 and is unrelated to 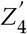.

#### Brownian Motion on a Tree

The most commonly used process is the Brownian Motion (BM) (Felsenstein, 1985). Under this model, for a given continuous trait **Z**′ measured at the tips of the tree, the covariance between traits 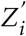 and 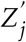 is simply proportional to the time of shared evolution between species *i* and *j*, i.e. the time *t_ij_* between the root of the tree and the most recent common ancestor of *i* and *j*: 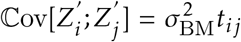, where 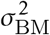 is the variance of the BM process. The expectation of each trait is equal to *μ*, the ancestral value of the process at the root.

#### Ornstein-Uhlenbeck on a Tree

To model stabilizing selection, the Ornstein-Uhlenbeck (OU) process is often used (Hansen and Martins, 1996; Hansen, 1997). Compared to the BM, it has an equilibrium value *β*, that represents the “optimal value” of the trait in a given environment. The trait is attracted to this optimum with a speed that is controlled by the selection strength *α*, or better the phylogenetic half-life *t*_1/2_ = log(2)*/α* (Hansen, 1997): when *t*_1/2_ is large compared to the total height of the tree *t* (*t*_1/2_ ≫ *t*), the trait needs a relatively long time to approach its optimum, and the selection strength is weak, while when *t*_1/2_ is small compared to *t* (*t*_1/2_ ≪ *t*), the selection strength is considered as strong. This process induces a different correlation structure than the Brownian motion, with stronger selection strength inducing weaker inter-species correlations (Hansen, 1997; Ho and Ané, 2013). Specifically, conditionally on a fixed root, 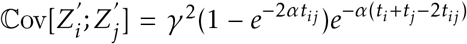, with 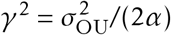 the stationary variance of the process, and *t*_i_ = *t_ii_* the time between the root and node *i* (Ho and Ané, 2013).

#### Within-Species Variation

The traditional PCM framework assumes that only one measurement is available for each species, and that there is no measurement error, i.e. that all the observed variation can be explained by the evolution process on the tree. However, ignoring measurement error can lead to severe biases (Silvestro et al., 2015; Cooper et al., 2016). In addition, in an inter-species RNA-Seq differential analysis, it is usual to have access to replicated measurements, i.e. to measurements for several individuals of the same species. There is a vast literature on the subject of within-species variation (Grafen, 1989, 1992; Lynch, 1991; Housworth et al., 2004; Ives et al., 2007; Hadfield and Nakagawa, 2010; Goolsby et al., 2017). One simple way to look at the problem in a univariate setting is to assume that all the individuals from a same species are placed on the tree as tips linked to a same species node with a branch of length zero (Felsenstein, 2008) and to add a uniform Gaussian individual variance *s*^2^ to all the tip samples traits (see Figures 1 and 2). In such a framework, the total variance of a sample trait *Z_i_* attached to a latent tip with trait 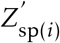 is given by 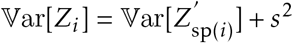, where 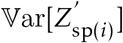 is determined by the chosen stochastic process to model the latent trait (BM or OU). Similarly, the covariance between two sample traits *Z_i_* and *Z_j_* attached, respectively, to latent tip traits 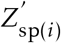 and 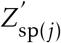 is given by 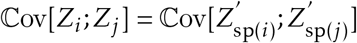.

**Figure 2:**
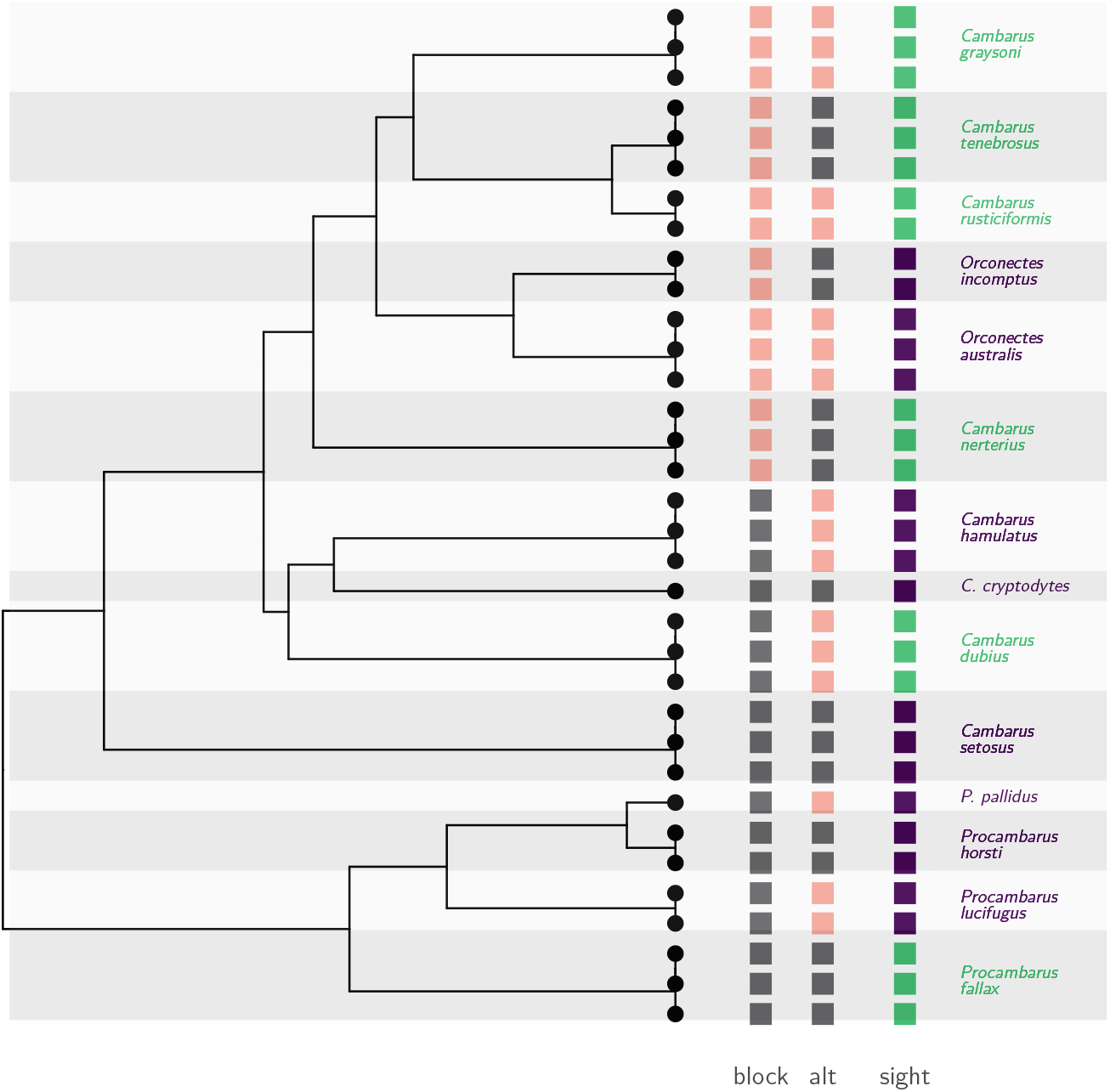
Time-calibrated phylogenetic tree of 8 blind (dark purple) and 6 sighted (light green) crayfish species (Stern et al., 2017). The root was dated to 65 million years before the present (Stern et al., 2017), but the tree was re-scaled to unit height for the analyses. The “sight” design (dark purple and light green squares) matches with the biological vision status of the species studied (Stern and Crandall, 2018a). The “block” and “alt” designs (light pink and gray squares) are artificial extreme scenarios representing, respectively, a situation where the design is almost un-distinguishable from the phylogeny-induced grouping (“block”), and a situation where groups are distributed evenly on the tree to maximize the contrast between sister species (“alt”).

### 2.6 Strategy 3: Phylogenetic Regression on Normalized Data

One way to include the phylogenetic structure with within-species variation, in statistical analyses is to use a Phylogenetic Mixed Model (PMM (Grafen, 1989, 1992; Lynch, 1991; Housworth et al., 2004)), where the vector 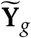 of the *n* normalized and transformed measurement for a given gene *g* is seen as the sum of a fixed effect, a random phylogenetic effect, and a random independent effect:

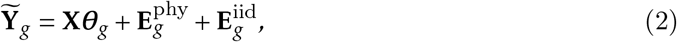

with **X** and ***θ***_*g*_ the design matrix and associated vector of coefficients as in Eq. (1), 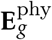 a vector of phylogenetically correlated residuals, with correlations given by the chosen process on the tree (see above) and 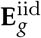 independent and identically distributed (iid) residuals, that can capture any non-phylogenetic source of variation of the data, such as within-species variation as described above.

Several methods for gene expression analysis based on models related to the PCM framework have been described in the literature, with different versions of the BM or the OU process, and with or without within-species variation (Khaitovich et al., 2004; Gu, 2004; Gu and Su, 2007; Bedford and Hartl, 2009; Rohlfs et al., 2014; Rohlfs and Nielsen, 2015; Gu et al., 2019), and in particular have been used to detect differences in gene expression across species (Brawand et al., 2011; Rohlfs et al., 2014; Rohlfs and Nielsen, 2015; Stern and Crandall, 2018a; Catalán et al., 2019; Chen et al., 2019).

For differential expression analysis, the *phylogenetic ANOVA* framework (Garland et al., 1993; Grafen, 1989; Rohlfs and Nielsen, 2015; Bastide et al., 2018) is particularly relevant, and can just be seen as the phylogenetic regression above, with the design matrix **X** encoding groups of species. This framework is for instance implemented in the popular and computationally efficient `R` package `phylolm` (Ho and Ané, 2014a).

## 3 Probabilistic Models and Data Simulation

Building on existing RNA-Seq methods (Robles et al., 2012; Soneson and Delorenzi, 2013; Soneson, 2014), we developed a new inter-species simulation framework that can generate realistic count datasets, and takes into account, first, the gene expression correlations induced by the phylogeny and, second, the different lengths a given gene can have in different species.

### 3.1 Realistic Simulations using the Negative Binomial Distribution

We briefly recall here the simulation framework detailed in (Soneson and Delorenzi, 2013), and implemented in `compcodeR` (Soneson, 2014).

#### Negative Binomial Distribution

Let *Y_gi_* be the random variable representing the count for gene *g* (1 ≤ *g* ≤ *p*) in sample *i* (1 ≤ *i* ≤ *n*), with true expression level *λ_gi_* and sampling depth *M_i_*. Following Robinson and Oshlack (2010), we model each count independently by a Negative Binomial (NB) distribution with expectation *μ_gi_* and dispersion *α_g_*, such that *Y_gi_* ~ `NB`(*μ_gi_*, *α_g_*) with:

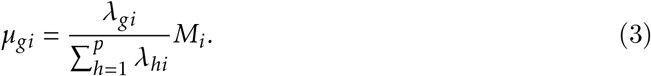

#### Differential Expression

To model differential expression, we assume that the samples are partitioned into two groups *S*_1_ and *S*_2_. For each gene *g*, the dispersion parameter *α_g_* is the same for all samples, while the expression level *λ_gi_* can only take two values: *λ*_*gS*_1__ if *i* is in *S*_1_ and *λ*_*gS*_2__ if *i* is in *S*_2_. Given *λ*_*gS*_1__, we take *λ*_*gS*_2__ as:

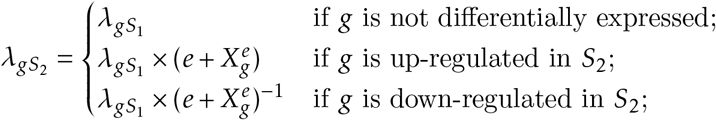

with *e* the minimal differential effect size, and 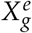 random variables independent identically distributed according to an exponential distribution with parameter 1. The values of the parameters are set to match the empirical counts expectation and dispersion of a real datasets.

### 3.2 Realistic Simulations using the Poisson Log-Normal Distribution

The Poisson Log-Normal (PLN) distribution has been advocated as an alternative to the NB distribution for the analysis of RNA-Seq data. Being more flexible, it is particularly well suited in the presence of correlations (Gallopin et al., 2013; Zhang et al., 2015; Choi et al., 2017), which proves essential for inter-specific datasets, as demonstrated in the next section. We show here how the parameters of a PLN model can be chosen to match first and second order moments of the NB model described above, making it possible to simulate realistic datasets under this more flexible framework.

#### The PLN Distribution

Under the PLN model, for each gene *g* and sample *i*, we assume that the observed count random variable *Y_gi_* follows a Poisson distribution, with log parameter a Gaussian latent variable *Z_gi_*, such that:

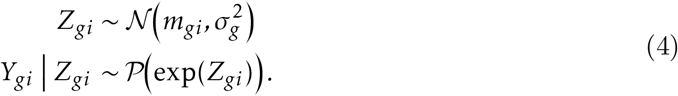

This model is similar in spirit to the NB distribution, that can be seen as Gamma-Poisson mixture (see e.g. Holmes and Huber, 2019, Chap. 4). Note that in both models the coefficient of variation of the mixing distribution is constant across samples for a given gene (Chen et al., 2014).

#### Matching Moments

Using standard moments expressions for the NB (Holmes and Huber, 2019) and PLN (Aitchison and Ho, 1989) distributions, it is straightforward to show that a PLN distribution with parameters *m_gi_* and 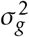 yields the same first and second order moments as a NB distribution with expectation *μ_gi_* and dispersion *α_g_* if and only if:

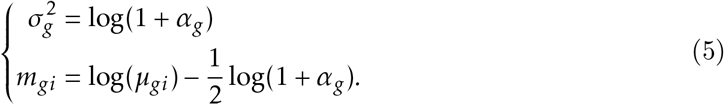

These equations allow us to readily use the framework developed in the previous section also in the case of a PLN simulation.

### 3.3 Taking the Phylogeny into Account with the Phylogenetic Poisson Log-Normal Distribution

In an inter-specific framework, various samples come from various species, which implies a specific correlation between measures, that can be taken into account in a multivariate PLN model, as shown below.

#### Continuous Trait Evolution Model

The models of trait evolution used in PCMs and presented in the previous section are generative, and can be used to simulate continuous traits at the tips of a tree (with possible replicates) such that their correlation structure is consistent with their phylogeny (see Fig. 1). Using a simple uniform Gaussian individual variance 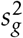 to model within-species variation, the trait variance **Σ**_*g*_ for the vector **Z**_*g*_ of continuous traits at the tips of the tree generated by such a process can be expressed as:

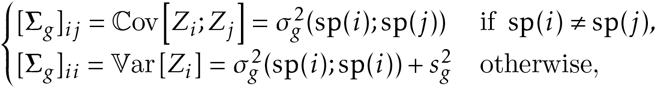

where 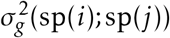 is the phylogenetic variance between species `sp`(*i*) and `sp`(*j*) of samples *i* and *j* (see Fig. 1), with a structure given by the evolution process (BM or OU, see expressions above), and 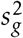 the added intra-species variation.

#### The Phylogenetic Poisson Log-Normal Distribution

The models described above are well suited for quantitative traits, but need to be adapted for count measures, such as the one produced by a RNA-Seq analysis. To handle such counts, we propose to add a Poisson layer to the trait evolution models described above, defining a “phylogenetic” Poisson Log-Normal (pPLN) distribution. More specifically, for a given gene *g*, we simulate a vector of *n* latent traits **Z**_*g*_ as the result of such a process running on the tree, and then, conditionally on this vector, draw the observed counts *Y_gi_* from a Poisson distribution with parameter exp(*Z_gi_*):

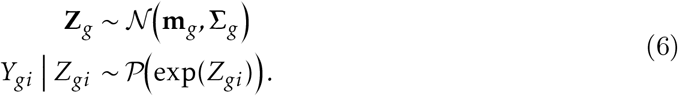

In other words, the vector of counts **Y**_*g*_ for each gene is drawn from a multivariate Poisson Log-Normal distribution, with parameters **m**_*g*_ and **Σ**_*g*_ obtained from the evolutionary models described above, **Σ**_*g*_ being the structured variance matrix of both phylogenetic and independent effects, and **m**_*g*_ a vector or expectations values at the tips, that can be set independently from the process.

#### Matching Moments for Realistic Simulations

Assuming that the diagonal coefficients of **Σ**_*g*_ are all equal to a single value 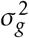, Equation (5) can be used to ensure that the pPLN model above yields the same marginal expectation and variance as a NB model with expectation *μ_gi_* and dispersion *α_g_*. At a macro-evolutionary scale, most of the dated phylogenetic trees encountered are ultrametric, i.e. are such that all the tips are at the same distance *t* from the root. In that case, all the phylogenetic models described above verify this variance homogeneity assumption. For instance, for the simple BM model with an extra layer of independent variation, we have 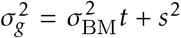. Note that, although the NB and pPLN models are set to have the same expectations and variance, they differ significantly in their covariances: while in the standard NB model, all the samples are independent from one another, in the proposed pPLN framework the measurements are correlated, with a structure reflecting both the tree and the selected evolutionary process.

### 3.4 Taking Differential Gene Lengths into Account

#### Length Normalisation of Counts

Let *ℓ_gi_* denote the length of the gene *g* for sample *i*. Following Robinson and Oshlack (2010), we take this length into account by changing Equation (3) to:

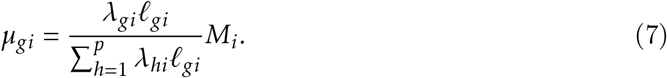

Note that the same overall sequencing depth *M_i_* is attributed to each sample, but that, because of the weighted average, it is preferentially allocated to longer genes.

#### Lengths Simulation

The lengths are simulated according to the pPLN model described above, with expectations and dispersions empirically estimated from the dataset at hand.

## 4 Simulation Studies

### 4.1 Material and Methods

#### Gene Expression Underlying Vision Loss in Cave Animals

We used our new simulation framework to generate realistic synthetic datasets, that were set to mimic the features of a recently published inter-species RNA-Seq dataset (Stern and Crandall, 2018a), while varying the level of evolutionary dependence. In this study (Stern and Crandall, 2018a), the authors analyzed the molecular mechanisms involved in vision loss in the North American family *Cambaridae* of crayfish species. They selected 8 blind and 6 sighted crayfish species, for which a time-calibrated maximum likelihood phylogeny is known (Stern et al., 2017). 3560 orthologous gene expressions were estimated using the method RNA-Seq by Expectation Maximization (RSEM) (Li and Dewey, 2011), with one to three replicates per species (see Fig. 2).

#### Base Simulation Parameters

Following the methodology described in the previous section, we simulated a “base scenario” dataset using the estimated crayfish tree re-scaled to unit height (*t* = 1), with the observed vision status design (“sight” design, see Fig. 2), and matching the empirical counts and gene lengths expectation and dispersion. The expression level *λ*_*gS*_1__ and the dispersion *α_g_* were estimated from the dataset for each gene *g*, while for each sample *i* the simulation sequencing depth *M_i_* was independently drawn from a uniform distribution with bounds *M*_min_ and *M*_max_ the observed empirical minimal and maximal values of the library size across all samples. We used a BM model of trait evolution, with an independent layer of individual variation 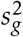 representing 20% of the total tip variance 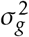 for each gene *g*: 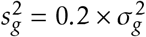, with 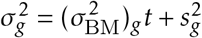. We chose a base effect size of 3, with 150 differentially expressed genes out of the 3560 simulated ones. From this base scenario, we varied several parameters in order to study their impacts on the simulated data. Each scenario was replicated 50 times.

#### Star Tree and NB Simulations

To check that our new pPLN framework produced datasets with properties similar to the well known NB framework, we replaced the crayfish tree with a star-tree, that mimics the NB situation where all species and replicates are independent.

#### Tree Group Design

The group design on the tree is known to strongly impact the properties of the data, in particular through its “phylogenetic effective sample size” (Ané, 2008; Bartoszek, 2016). To study its effect in a gene expression context, we replaced the “sight” design with a “block” and “alt” design (see Fig. 2), that were chosen to model two extreme situations. In the “block” design, all the species with a given group are nested within a single clade, so that the differential expression signal is redundant with the phylogenetic signal. At the other end of the spectrum, the “alt” design was chosen so that sister species are in different groups, in order to maximize the contrast between organisms that share a long common history. We expect the “alt” design to produce datasets with a stronger signal.

#### Differential Analysis Phylogenetic Asymptotic Effective Sample Size

To quantify the intrinsic difficulty of a design compared to another, we propose a new “differential analysis phylogenetic asymptotic effective sample size” (dapaESS). Given a phylogenetic tree 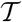, we first remove all replicates, so that there are no zero-length branches. Then, given a design vector **x**, we postulate a simple BM model for an hypothetical continuous trait **y** at the tips: **y** = *θ*_0_**1** + *θ*_1_**x** + *σ***e**^*BM*^, with 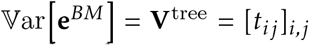. From standard linear model theory, the variance of the maximum likelihood estimator of the coefficient *θ*_1_ is given by (Ané, 2008): 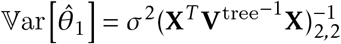, with **X** = (**1 x**) the matrix of predictors. We hence define: 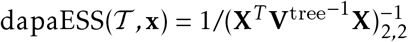. In the case where all the species are independent (star-tree 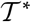), we fall back on a standard differential expression analysis, and we get, assuming that there are *n* species and that the groups are balanced: 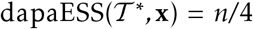, which is the standard effective sample size for a balanced two-sample t-test with uniform variance. This gives us a base-line for a “standard” difficulty, and we use in the following the normalized dapaESS: 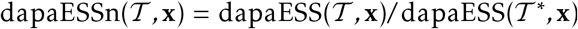. A value lower than 1 indicates a design that is deemed more difficult than a standard independent design (larger asymptotic variance of the estimator), while a value greater that 1 indicates a problem where the phylogeny actually helps in finding the significant differences. Note that this score can be computed *a priori*, and, as shown below, can be used to asses the quality of the experimental design.

#### Simulation Process

The simulation process impacts the tree induced correlation between species (Blomberg et al., 2003; Harmon, 2019). To study the impacts of this modeling choice, we replaced the BM process with an OU, with a phylogenetic half-life (Hansen, 1997) *t*_1/2_ = log(2)*/α* fixed equal to 50% of the tree height.

#### Within-Species Variation Level

We mitigated the effect of the BM model on the tree by varying the level of the independent individual variation representing 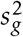, from 40% to 0% (i.e., all the measurements from a same species are perfectly correlated).

#### Inference Methods Used

We chose the following statistical inference methods, representing the three main approaches presented above: `DESeq2` (Love et al., 2014) assumes a NB distribution on independent counts; `limma` (Ritchie et al., 2015) applies an Empirical Bayes moderation (without a mean-variance trend correction, unless otherwise specified) on independent normalized counts, possibly assuming that all the samples in a same species are correlated (`limma cor` (Smyth et al., 2005)); and `phylolm` (Ho and Ané, 2014a) uses a phylogenetic regression framework based on a BM or OU process, with measurement error. For `phylolm`, the differential analysis relied on a t statistic computed for each gene independently, conditionally on the estimated maximum likelihood parameters (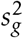 and *α_g_* for the OU). The raw p-values computed by all methods were adjusted using the `BH` method (Benjamini and Hochberg, 1995), using the `R` function `p.adjust`. Inferred gene expression differences across groups were marked as significant if their associated adjusted p-value was below the threshold of 0.05.

#### Length Normalisation and Transformation

In `DESeq2` (Love et al., 2014), we used the default RLE method (Anders and Huber, 2010) to compute the sample-specific normalization factor *m_i_*. We followed the recommendations of the section “Sample-/gene-dependent normalization factors” from the `DESeq2` vignette to compute the coefficients *c_gi_* from the coefficients *m_i_* and gene lengths *ℓ_gi_* detailed in section 2. For methods requiring a preprocessing normalization of the count data (`limma` and `phylolm`), we used the TMM method (Robinson and Oshlack, 2010) implemented in the `calcNormFactor` function in `edgeR`, and a TPM length normalization with a log_2_ transformation. We studied the effect of these choices by testing combinations of other normalization methods (RPKM length normalization; or a simple CPM, i.e. no length normalization), and an other transformation function (square root, as advocated in Musser and Wagner (2015)).

#### Scores Used

To assess the performance of the inference methods, based on the list of true (simulated) differentially expressed genes, we computed the number of True Positives (TP), True Negatives (TN), False Positives (FP) and False Negatives (FN). We used the Matthews correlation coefficient (MCC = [TP·TN–FP·FN]·[(TP+FP)(TP+FN)(TN+FP)(TN+FN)]^-1/2^) as advised in Chicco and Jurman (2020). We also computed the True Positive Rate (TPR = TP/(TP + FN)) and the False Discovery Rate (FDR = FP/(FP + TP)). In addition, we compared the features of the simulated datasets with the empirical one using the `countsimQC R` package (Soneson and Robinson, 2018).

### 4.2 Results

#### PLN and NB Simulation Frameworks Produce Similar Datasets

When parametrized to produce the same moments, the pPLN framework on a star tree produces datasets that are similar in difficulty to the classical NB framework (Fig. 3, first two columns). While `limma` controls the FDR to the nominal rate, `DESeq2` fails to control the FDR in this case with a lot of variance (empirical dispersion range from 0.1 to 5, see also Fig. 5). As it assumes a NB distribution of the counts, `DESeq2` suffers from the deviation from this model, as opposed to `limma`, which performs equally well in both cases. As showed by the `countsimQC` analysis, the datasets simulated with the pPLN and the NB frameworks have similar features, and are comparable to the original empirical dataset (data not shown, comparison report available on the GitHub repository github.com/i2bc/InterspeciesDE). While preserving the univariate moments, the tree included in the framework (Fig. 3, last column), introduces some phylogenetic correlations between the species, and leads to a spectacular loss of power of both methods, with a rate of false discoveries higher than three quarters.

**Figure 3:**
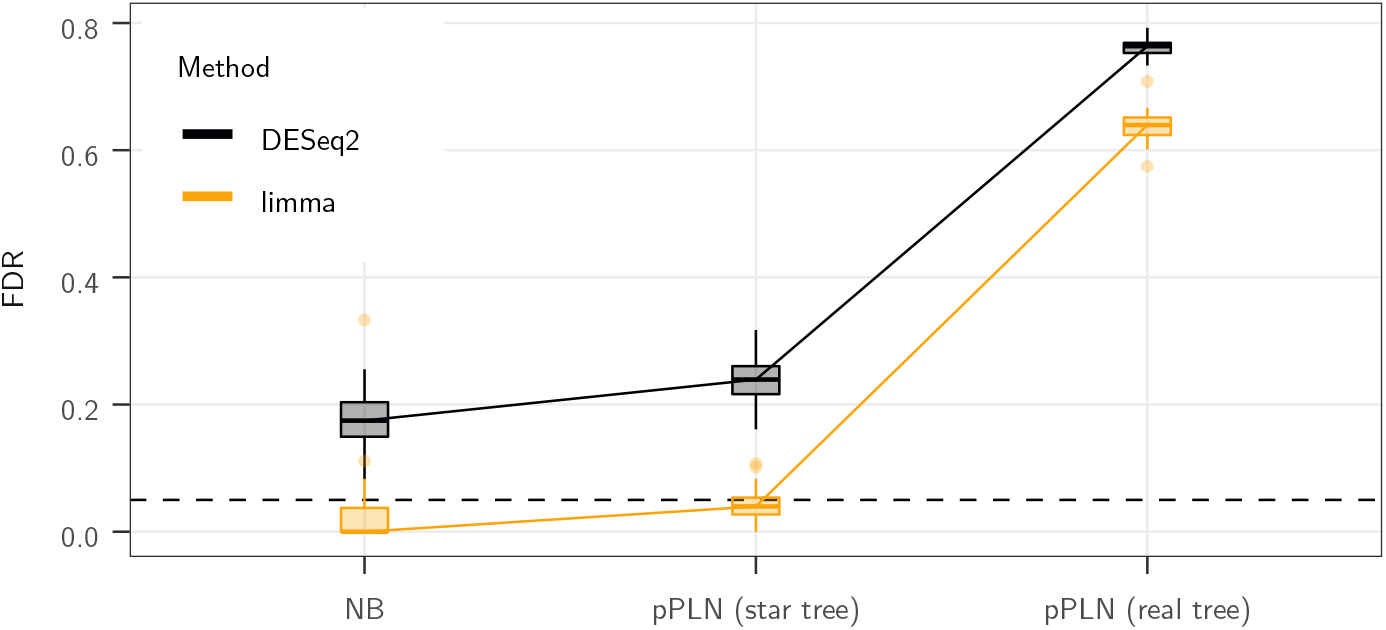
The base scenario (`pPLN` (`real tree`), right) has empirical moments drawn from (Stern and Crandall, 2018a), with an effect size of 3, a BM model of evolution with added intra-species variation accounting for 20% of the total variance, on the maximum likelihood tree, with the observed `sight` groups (see Fig. 2). It is compared to a pPLN model with the same parameters, but in a case where all samples are independent (`pPLN` (`star tree`), middle), and to a NB model with the same moments and effect size (`NB`, left). The `DESeq2` (black) and `limma` (light orange) inference methods are applied to each scenarios, and their FDR is compared. The black dashed line represents the nominal rate of 5% used to call positives. For `limma`, the counts are normalized using log_2_(TPM) values. Boxplots are on 50 replicates.

#### Phylogenetic Data Requires Correlation Modeling

For data simulated according to the base scenario, methods that explicitly model sample correlations (`limma cor` and `phylolm`) perform best (Fig. 4, dark purple line). `limma cor` exhibits the best behavior with the highest MCC, and a TPR reaching about 80%. Its FDR is still above the nominal rate (median around 10%).

**Figure 4:**
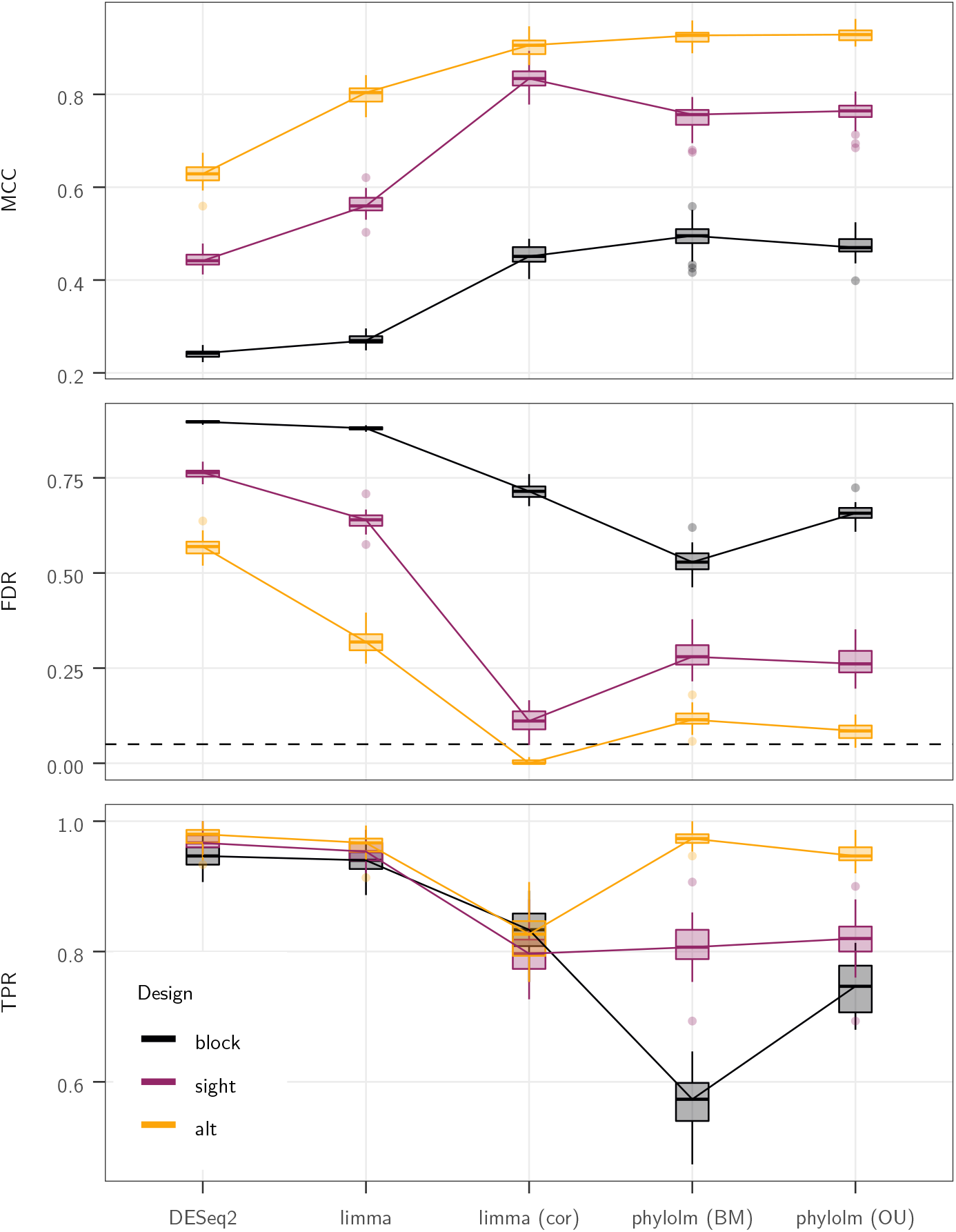
Results in term of MCC (top), FDR (middle) and TPR (bottom) scores of the five selected statistical methods (x axis) on the pPLN base scenario, that has an effect size of 3, a BM model of evolution with added intra-species variation accounting for 20% of the total variance, on the maximum likelihood tree (Stern and Crandall, 2018a), with the observed `sight` groups (dark purple line, see Fig. 2). The `alt` (light orange line) and `block` (black line) groups are also tested, with the same parameters. For the FDR, the black dashed line represents the nominal rate of 5% used to call positives. When required, the counts are normalized using log_2_(TPM) values. Boxplots are on 50 replicates.

#### Tree group Design Matters

The `alt` designs produces datasets with the clearest signal, (Fig. 4, light orange line). In this case, `limma cor` is able to correctly control for the FDR. Although `phylolm` methods have slightly higher FDR, they achieve a better TPR reaching almost 100%, leading to a better overall MCC score. At the opposite of the spectrum, the `block` designs produces datasets with a very weak signal, with differentially expressed genes counts M-A values strongly overlapping a very diffuse non-differentially expressed genes distribution (Fig. 5). All methods applied to the `block` design have FDR higher or equal to about 50% (Fig. 4, black line). The BM `phylolm` tool has the least bad MCC score (about 0.5), although with the worst TPR (around 50%). The relative difficulties of each design is correctly captured by the normalized dapaESS. While the `block` design has a lower dapaESS than the independent case (dapaESSn = 0.69), the `alt` design has a higher one (5.1), and the `sight` design lies in the middle (1.4).

**Figure 5:**
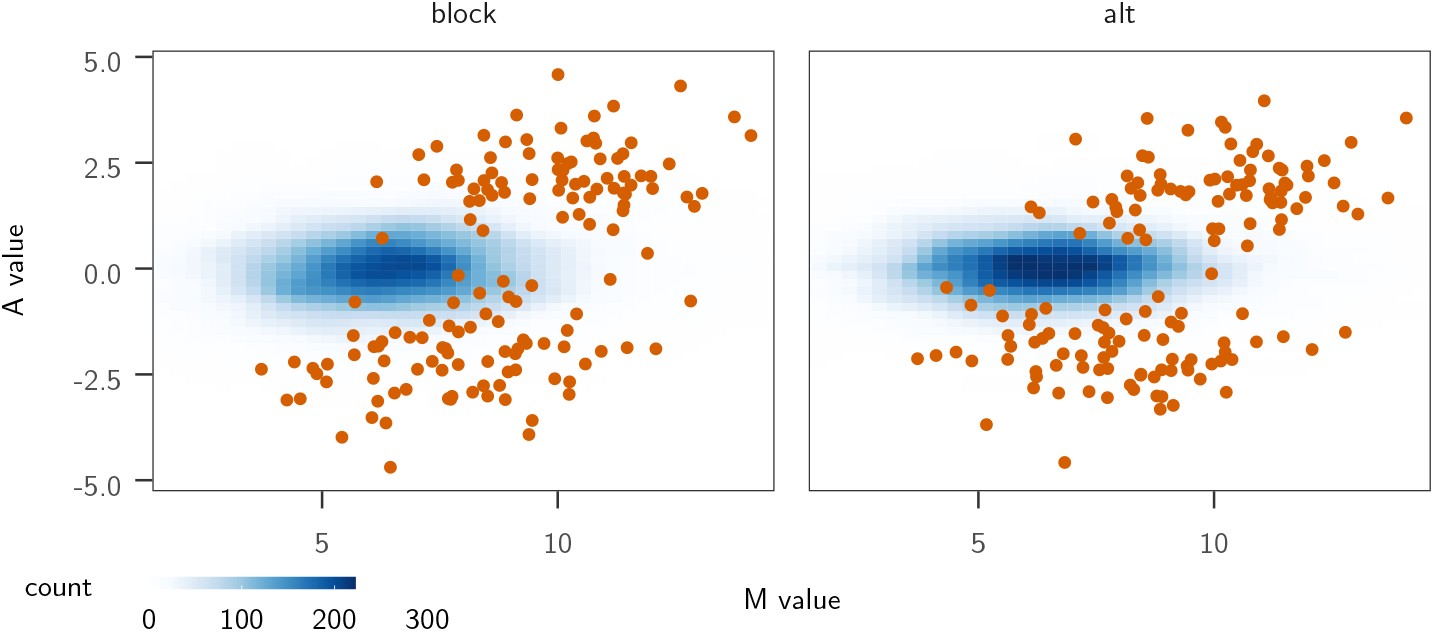
M-A plots (log2 fold change as a function of the mean of normalized counts for each gene) of the datasets produced with base pPLN parameters (effect size of 3, BM model with added intraspecies variation accounting for 20% of the total variance), on the maximum likelihood tree (Stern and Crandall, 2018a), with the `block` (left) and `alt` (right) designs. The M-A values distribution for the 3410 non-differentially expressed genes is shown as a tile plot, with deeper blues representing high probability values. The M-A values of the 150 differentially expressed genes are shown as red dots.

#### OU Makes the Signal Weaker and is Hard to Correct For

When simulating the counts using an OU model of trait evolution for the latent trait instead of a BM, the signal becomes weaker, and all methods achieve lower MCC scores (Fig. 6). The `limma cor` methods performs the best in this case, even when compared to a `phylolm` method that explicitly takes the OU model into account.

**Figure 6:**
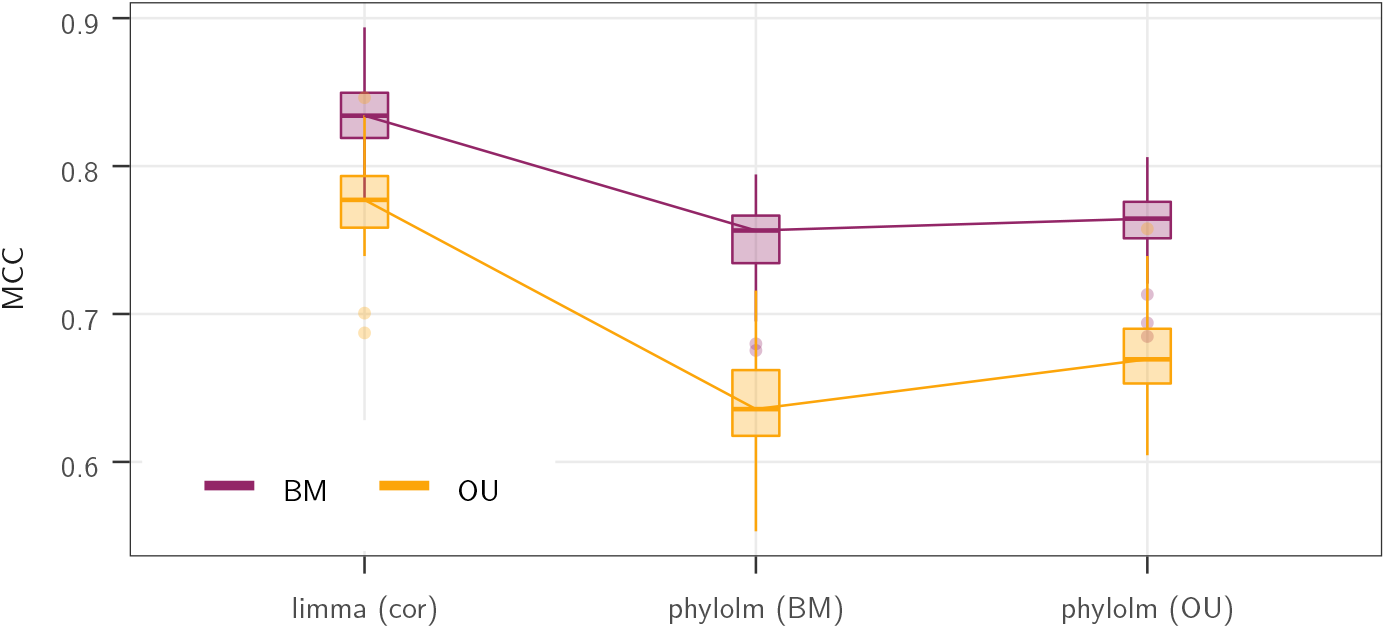
Results in term of MCC scores of the three correlation aware statistical methods (x axis) on the pPLN base scenario (effect size of 3, intra-species variation accounting for 20% of the total variance), with a BM (dark purple line) or an OU (light orange line) model of evolution on the maximum likelihood tree (Stern and Crandall, 2018a), with the observed `sight` groups (see Fig. 2). The counts are normalized using log_2_(TPM) values. Boxplots are on 50 replicates.

#### Phylogenetic Methods are Robust to Intra-Species Variations

When reducing the intraspecific variance to 0 (inducing a correlation of 1 between sample values of the same species), the `limma cor` method loses its advantage compared to the `phylolm` methods, which performances are less affected by the level of intra-specific noise (Fig. 7).

**Figure 7:**
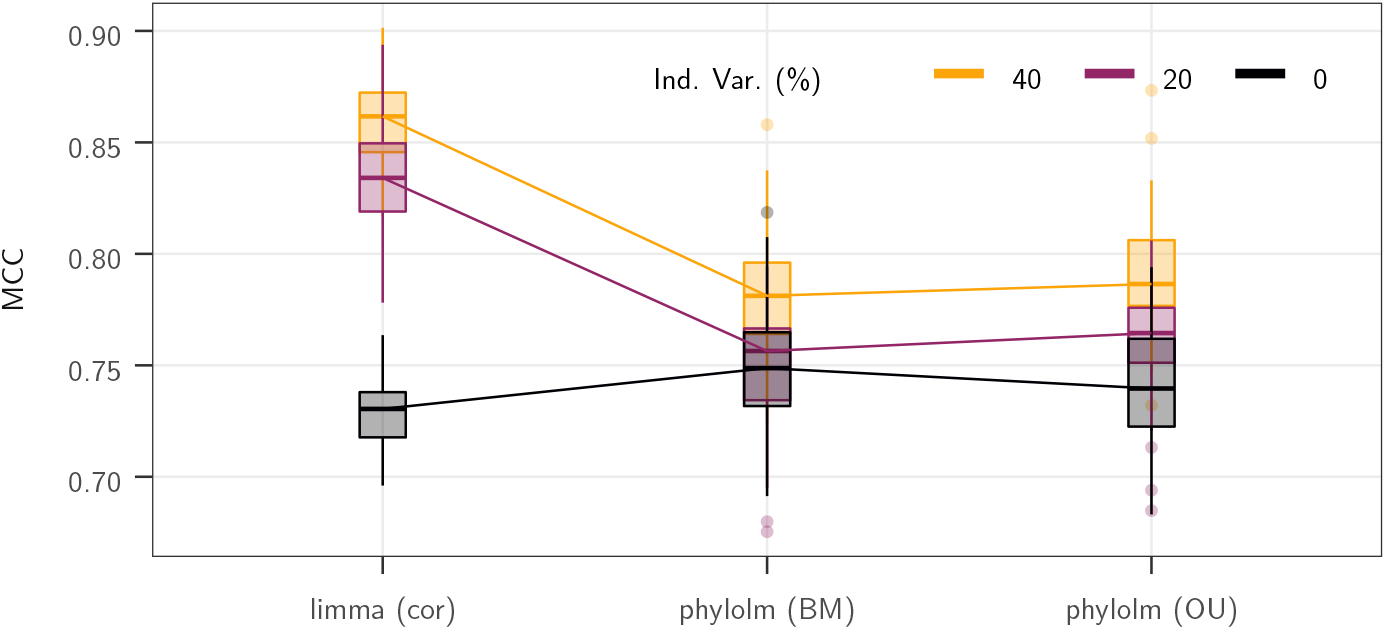
Results in term of MCC scores of the three correlation aware statistical methods (x axis) on the pPLN base scenario with an effect size of 3, a BM model of evolution on the maximum likelihood tree (Stern and Crandall, 2018a), with the observed `sight` groups, and intra-species variation accounting for 40% (light orange line), 20% (dark purple line), or 0% (black line) of the total variance). The counts are normalized using log_2_(TPM) values. Boxplots are on 50 replicates.

#### log_2_(TPM) Normalisation is Slightly Better on Phylogenetic Data

Taking gene lengths into account, using either TPM or RPKM, significantly improves the power of the methods, in particular in term of TPR (Fig. 8). Although TPM normalization leads to a slightly better MCC median, its performances are largely similar to the RPKM normalization. On this base scenario, the log_2_ transformation leads to a consistent gain of about 10% in TPR compared to the square root (going from around 70% to 80%, Fig. 8).

**Figure 8:**
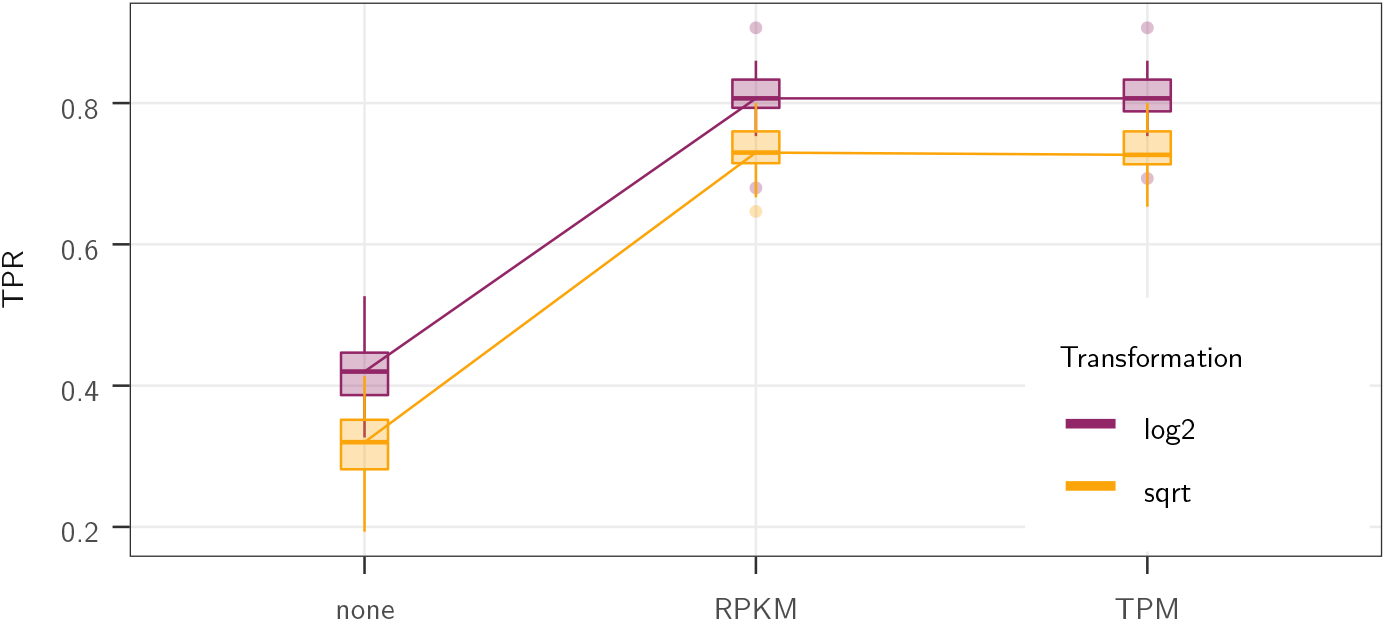
Results in term of TPR score of the `phylolm` (`BM`) method on the pPLN base scenario (effect size of 3, BM model of evolution on the maximum likelihood tree (Stern and Crandall, 2018a), with the observed `sight` groups, and added intra-species variation accounting for 20% of the total variance). The counts are length-normalized (x axis) using CPM (length not taken into account, `none`), RPKM or TPM, and transformed using the square root (light orange) or the log_2_ (dark purple) functions. Boxplots are on 50 replicates.

#### No Small Counts in De Novo Assembled Data

Including a mean-variance trend correction in the `limma cor` method did not change its performance on the base scenario, producing very similar MCC values (the median MCC on all 50 runs differ by less than 0.002). This is consistent with the fact that the original dataset uses *de novo* assembled data, that naturally exclude any small counts, and hence the need for a mean-variance trend correction (see Discussion).

## 5 Discussion and Conclusion

### 5.1 Simulation Study

Our targeted simulation study illustrates some of the specificities of inter-species RNA-Seq differential expression analysis. First, it is essential to take the correlation between replicates within a given species into account. Failure to do so leads to very high rates of false discoveries (Fig. 4), that make the analysis unreliable and hard to exploit. Indeed, the `limma` method with added correlation seems to outperform other tools, including phylogenetic comparative methods, in many settings. These results tend to indicate that, even if the full tree is not included in the analysis, incorporating these simple correlations between replicates might be sufficient to efficiently analyse inter-species datasets, at least for some simulation designs. The group design on the tree was indeed found to be extremely important (Fig. 4). A balanced design, were the groups are evenly spread over all clades, has a stronger signal (Fig. 5), and allows the analysis to be abstracted from the phylogeny to some extent, as classical tools for differential expression analysis work best in this configuration. On the other hand, when the groups are clustered in the phylogeny, the signal is weaker as it becomes more difficult to distinguish the real group effect from the simple drift that tends to isolate clades from one another. This is in particular the case of designs where one clade or species is tested against out-groups, that is sometimes encountered in the literature (Brawand et al., 2011; Rohlfs and Nielsen, 2015). In this configuration, phylogenetic comparative methods, although imperfect, are essential. Finally, this study confirms the importance of length normalisation for inter-species differential gene expression analysis to achieve acceptable power detection levels (Fig. 8). Although we did not find any significant difference in performance between RPKM and TPM normalizations, the log_2_ transformation seemed to have a slight advantage over the square root in this simulation setting.

### 5.2 Simulation Design

In this work, we proposed a method to simulate RNA-Seq gene expression across multiple species. Similar to intra-species simulation tools (Dillies et al., 2013; Soneson and Delorenzi, 2013; Soneson, 2014), our simulation method can use empirical datasets to set the value of parameters such that the simulated datasets are as close as possible to the real ones, with matching empirical marginal expectation and variance. When applied to independent species, it produces datasets with comparable features (Fig. 2). In our specific simulation studies, we use the dataset from Stern and Crandall (2018a). This dataset was obtained using *de novo* assembled data. In addition, we focused on genes with one-to-one orthologuous relationships across species. As a consequence, this dataset had a low number of zeros and small counts, and a large variance across samples. The simulated datasets had similar characteristics, which could explain the low performance of `DESeq2`, even when the data was simulated without correlation (Fig. 2), and the fact that the trend procedure did not add any power to the `limma` method. Inter-species RNA-Seq gene expression datasets are very diverse, with specificities depending on the underlying biological question being studied. This work provides a first step toward realistic simulation of such datasets.

### 5.3 Simulation Tool

Compared to classical intra-species simulation tools (Dillies et al., 2013; Soneson and Delorenzi, 2013; Soneson, 2014), our simulation framework incorporates the species tree and the gene length, which may vary across species. It makes it possible to model the evolution of gene expression on the tree using two different processes (BM or OU), and it allows for additional independent variation, that can model e.g. inter-specific variation or measurement error. This complex model leads to new effects, that can be difficult to predict. In particular, we showed that the distribution of the groups on the tree had strong effects on the ability of all methods to detect a group expression shift. We proposed a normalized criterion (dapaESS) to assess the difficulty of the group design for the differential gene expression analysis problem. Although it does not take into account the number of replicates or the specific evolution model, we showed that it could well represent the difficulty of an experimental design. The strength of this criterion is that it only depends on the timed species tree and the tips group allocation, and can be computed before any statistical inference or even data collection. It can hence be used as a practical guide on the expected power of the experimental design. In this review, we focused our attention of the detection of shifts of expression between groups spanning across species. However, inter-species datasets are also used to address many other questions, such as equality of within-species variance, expression divergence, or detection of neutral versus directed evolution regimes. Several tools from the PCM literature have been used to this end, that rely on various models of trait evolution with appropriate parameter constraints. Since our simulation tool is modular, those various processes could be implemented, in order to produce realistic RNA-Seq datasets with the desired structure. Such an extended framework could help researchers to test the statistical properties of these complex inference models.

### 5.4 Inference Tools

In this study, we focused on a few inference tools, that come either from the RNA-Seq or the PCM literature, limiting ourselves to methods implemented in `R` and that can do differential analysis. Although a more comprehensive simulation design would be needed to draw stronger conclusions, our results show that simulations under the OU model lead to more difficult datasets, and that even methods that include the OU model in their framework fail to completely correct for this effect. This could be linked with the fact that the estimation of the selection strength in an OU model is a notoriously difficult question, especially on an ultrametric tree (Ho and Ané, 2014b; Cooper et al., 2016). Having to estimate this parameter for thousands of genes is bound to generate some instability, and to deteriorate the performance of those tools. Gu et al. (2019) recently proposed an empirical Bayes approach to deal with this parameter in an RNA-Seq setting. One possible direction could be to adapt this method to a differential analysis problem. More generally, our simulation studies illustrate the need for new statistical tools for inter-species differential analysis, that would combine the strengths of both the classical RNA-Seq literature, that can deal with the specificities of this noisy data, and the PCM literature, that takes into account the phylogeny, an information that can be crucial to correctly interpret inter-species data.

## 6 Key Points

- Inter-species RNA-Seq datasets have a complex structure, and require a dedicated simulation tool that can generate count data with phylogeny induced correlations and that can take varying gene lengths into account.
- Differential analysis for inter-species RNA-Seq data requires a tool that can take at least within-species sample correlations into account and an adequate length normalisation procedure.
- The experimental design of the group allocation on the phylogeny has a strong impact on the differential expression signal, and is well captured by the dapaESS score, that can be computed *a priori* before any statistical analysis or data collection.

## 7 Data and Code Availability

The simulation tool is integrated into the `compcodeR` package, that is freely available on the Bioconductor platform, and documented through a specific vignette (doi.org/10.18129/B9.bioc.compcodeR). The data and code used for the simulation study are available on the following GitHub repository: github.com/i2bc/InterspeciesDE.

## 8 Acknowledgments

We thank David Stern for sharing his insights on the dataset, and for giving us access to the raw count data. P.B. and M.G are grateful to Sylvain Merlot for initiating this project, to Marie-Laure Martin and Guillem Rigaill for useful discussions, and to Claire Ducos, Marie Michel and Sarah Jelassi for their work during their master internship. This work was partly funded by the I2BC and the MI CNRS through the MODELCOG (M.G.) and X-TrEM projects (Sylvain Merlot). We are grateful to the INRAE MIGALE bioinformatics facility (MIGALE, INRAE, 2020. Migale bioinformatics Facility, doi: 10.15454/1.5572390655343293E12) for providing computing and storage resources.

